# The behaviour bends, the code holds: working-memory category information in the multiple-demand network across graded interruption

**DOI:** 10.64898/2026.11.03.763342

**Authors:** Olga Leticevscaia, Eva Feredoes

**Affiliations:** Centre for Integrative Neuroscience and Neurodynamics, Reading, University of Reading, United Kingdom; San Jose State University, San Jose, United States

## Abstract

Protecting working memory (WM) from interference is essential for goal-directed behaviour, yet its neural mechanisms remain unclear. The multiple-demand (MD) network — a set of frontoparietal regions recruited by diverse cognitive demands — is a candidate system for maintaining WM content under interference. We examined how the MD network represents a memorandum when an unrelated task interrupts maintenance. Participants memorised an image of a tool or animal and, during the delay, performed a digit task whose difficulty was parametrically varied (passive, easy, hard), allowing us to test how task engagement and associated cognitive demands affected concurrently maintained information.

Using fMRI and multivariate pattern analysis, we decoded the maintained category (tool vs. animal) from 13 MD regions and early visual cortex. Behaviourally, interruption reduced recall accuracy, while the most demanding interrupter slowed responses. Neurally, category information was present during the delay, most strongly in visual cortex and bilateral intraparietal sulcus. This information was unaffected by task engagement (active vs. passive), consistently across decoding accuracy and cross-validated Mahalanobis (crossnobis) distance, with Bayesian evidence supporting the null effect. Increasing the demand of the active interrupter, however, produced a modest, network-wide reduction in crossnobis distance that was not detectable in decoding accuracy.

Together, these findings reveal a dissociation between the behavioural costs of interruption and the neural representation of maintained category information. Although interruption impaired recall, engaging in an interrupter did not measurably alter category-level information in the MD network or visual cortex. Increasing the interrupter’s demand produced only a modest, network-wide reduction in representational distance, without affecting decoding accuracy, suggesting that the behavioural costs of interruption may not be fully explained by changes in the categorical representation of the memorandum.

## Introduction

Our ability to maintain and process information in mind in an ongoing manner, known as working memory (WM) (Baddeley, 1992), provides us with the ability to achieve immediate behavioural goals as well as subserving higher-order cognitive processes (Oberauer, 2009). Irrelevant information coming from external and internal sources poses a challenge to successful WM, disrupting the maintenance-related processes, resulting in forgetting (Kiyonaga & Egner, 2013; Lorenc et al., 2021). Understanding how WM contents are protected from interference is therefore central to understanding the mechanisms underlying memory maintenance.

One approach is to investigate the impact of distracting stimuli on WM representations being held in the brain. Recent fMRI decoding studies have examined how distractors influence WM representations (Bettencourt & Xu, 2016; Kiyonaga et al., 2017; Lorenc et al., 2018; Rademaker et al., 2019; Hallenbeck et al., 2021; Vu-Cheung et al., 2021). Reduced decoding accuracy of WM stimuli during distraction has been interpreted as disruption of the neural patterns supporting maintenance, potentially reflecting competition between memoranda and distractors for shared neural resources (Hallenbeck et al., 2021). Across these studies, the intraparietal sulcus (IPS) has been implicated in maintaining WM representations in the presence of visual distraction. Bettencourt and Xu (2016) found that visual distractors disrupted orientation decoding in occipital cortex but spared decoding in superior IPS, suggesting that IPS representations are less susceptible to sensory input. Lorenc et al. (2018) further showed that orientation information was decodable from IPS specifically when distractors were expected, suggesting that mnemonic representations in this region might be recruited adaptively according to task demands. Consistent with this account, Rademaker et al. (2019) found that distraction shifted memorised orientations in IPS from a sensory to a more abstract mnemonic format, whereas representations in the visual cortex remained predominantly sensory and more susceptible to distortion. Together, these findings suggest that IPS can support distraction-resistant WM representations, potentially by flexibly transforming memoranda into a more durable, interference-resistant format.

Other studies, however, suggest that mnemonic representations can remain resilient to interference even within the sensory cortex, as indicated by preserved decoding following intervening stimuli (Hallenbeck et al., 2021; Vu-Cheung et al., 2021). Hallenbeck et al. (2021) reliably decoded remembered polar angle from sensory, parietal and frontal regions despite intervening moving-dot stimuli, while Vu-Cheung et al. (2021) found no reduction in spatial-location decoding within retinotopic cortex following a response-demanding interrupter. Thus, mnemonic representations can remain stable across the sensory cortex despite intervening sensory input, potentially reflecting the presence of robust memory codes in these areas. However, whether such resilience extends to interruptions that shift attention and temporarily present new cognitive demands remains less understood. Unlike passive distractors, active interrupters require the participant to engage additional cognitive operations, raising the possibility that memory protection depends not only on stable posterior representations but also on the continuity of cognitive control.

Evidence for such control comes from studies of active distraction. Kiyonaga et al. (2017) found that decoding accuracy for memorised stimuli in sensory regions varied with the difficulty of an intervening visual-search task, suggesting competition for representational resources, with prefrontal cortex more engaged when both memory load and search difficulty were high. These findings implicate prefrontal control in resolving interference, but such control is unlikely to be reliant on the prefrontal cortex alone (Menon & D’Esposito, 2022). Indeed, Clapp et al. (2010) showed that a passively presented distractor left frontal–sensory connectivity intact, whereas a response-demanding interrupter disrupted this connectivity, which was restored only post-interruption, highlighting the cognitive control demands of dealing with an interruption during WM maintenance.

A candidate system for exerting such control is the multiple-demand (MD) network—a set of frontal and parietal regions, including the intraparietal sulcus (IPS), inferior frontal junction (IFJ), anterior and posterior inferior frontal sulcus (aIFS, pIFS), premotor cortex (PM), anterior insula/frontal operculum (AIFO), and pre-SMA/anterior cingulate cortex—that is engaged across WM, attention and other cognitively demanding tasks (Duncan, 2010; Fedorenko et al., 2013; Assem et al., 2020). Importantly, the MD network does not simply represent all available information; rather, it flexibly prioritises information according to current task demands. When attention is directed to one feature of a stimulus, MD regions preferentially represent the attended feature while excluding irrelevant information, even when both features are represented in early visual cortex (Jackson et al., 2017; Jackson & Woolgar, 2018). The strength of task-relevant information coding in MD network increases with task difficulty (Woolgar et al., 2011), with dlPFC playing a critical role in this enhancement of task-relevant representations (Jackson et al., 2021). Consistent with this prioritisation, MD regions have been shown to represent cognitive operations to a greater degree than incidental stimulus categories, which are represented more robustly in the sensory cortex (Shashidhara et al., 2024). Together, these findings characterise the MD network as an adaptive coding system that flexibly prioritises information according to its current relevance.

This raises a critical question for WM: how does the MD network represent a memorandum that remains relevant to the trial but is not relevant to an intervening task? To address this question, we asked participants to maintain a visual object in their WM during an fMRI scanning session. Each trial, a delay period task (interrupter) would appear and require participants’ response; interrupter demand was varied parametrically (passive, easy, hard) while the visual display was held relatively similar among conditions, aiming to isolate cognitive demand from perceptual interference. Critically, the memorandum remained relevant across the interrupter task, but was never relevant to its requirements. We used neural decoding and cross-validated representational distances (crossnobis) to track a maintained object category (animal or tool) across MD regions, for all interrupter conditions. We predicted that if the MD network actively protects task-relevant content, category information should be *enhanced* at higher levels of demand, consistent with the demand-dependent enhancement of task-relevant codes reported in this network (Woolgar et al., 2011; Jackson et al., 2021). If the interrupter instead competes for shared resources, category information should *decline* as demand increases. Finally, if MD regions preferentially represent information relevant to the task currently at hand, category information—relevant to the trial but incidental to the interrupter—should remain detectable but *unchanged* across demand (Jackson & Woolgar, 2018; Shashidhara et al., 2024). Thus, the relationship between category information and interrupter demand directly distinguishes three accounts of MD function: active protection, competitive interference, and demand-insensitive maintenance. These results would provide further evidence on the nature of MD network involvement in the representation of task relevant information across a variety of behavioural conditions.

## Methods

### Procedure

#### Participants

Twenty-eight participants (18 females; mean age=26.1, SD= 7.9) took part in the study. All participants had normal or corrected-to-normal vision and no history of neurological or psychiatric disorders. Participants gave written informed consent and were reimbursed for their time. The experiment was approved by the University of Reading Research Ethics Committee.

Data from two participants were excluded from the analysis due to low performance on the memory task (<60% recall accuracy). Since excessive movements may compromise the integrity of the fMRI analysis (Power et al., 2012; Van Dijk et al., 2012), after applying a 2 mm threshold for translation or rotation of the movement parameters, another four datasets were excluded after the realignment preprocessing step. Specifically, two participants had excessive movements in 3 runs, and another two, in 4 runs. The final sample resulted in twenty-two individual data sets with all runs included in the analysis.

Sample size of twenty-eight was chosen to be comparable to previous fMRI studies applying the decoding to the MD network (Woolgar et al., 2011; Jackson et al., 2017; Jackson & Woolgar, 2018). To characterise what the final sample could detect, we conducted a sensitivity analysis: with N = 22, a two-tailed paired t-test at α = .05 has 80% power to detect within-subject effects of dz ≥ 0.63 (90% power: dz ≥ 0.73). Smaller modulations of category information cannot be excluded by frequentist tests alone; we therefore complemented all null results with Bayes factors (see Bayesian analyses), which quantify the relative evidence for the null and alternative hypotheses given the observed data.

#### Memory Task Stimuli

The stimulus set for the memory task consisted of black-and-white morphed images of animals and tools. All images were generated using the diffeomorphic transformation algorithm (Stojanoski & Cusack , 2014), with an equal amount of transformation between each image. Each stimulus set for the training and for the main experiment consisted of one pair of animals and one pair of tools; three morphed exemplars were generated for each image. Figure 1, panel B depicts all the stimuli used as memory targets in the main experiment. The aim of this stimulus set was to make the memory task more difficult, because participants were required to memorise not just an object’s identity, but also its fine details, in order to select the correct item at recall.

**Figure 1.**
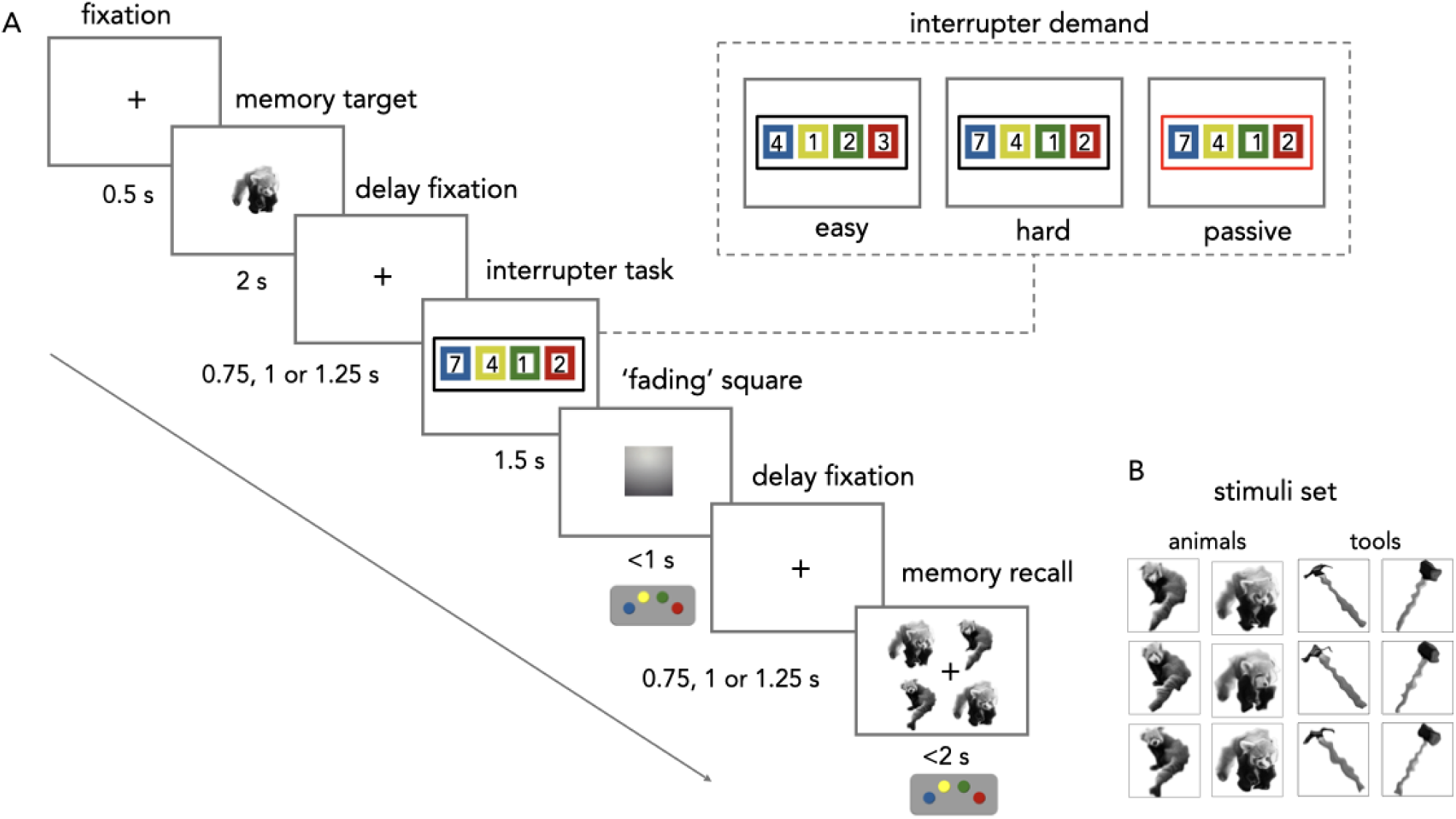
Dual Task Paradigm (A) and stimulus set for the memory task (B). (A) The dual-task paradigm involved a memory task and a digit task, with the latter serving as an interrupter during memory maintenance. Participants saw an image (tool or animal) to remember, followed by a fixation cross. Next, four coloured squares with digits appeared, and participants had to sort the digits and press the button to indicate the second item in the list before a ‘fading square’ disappeared. The frame colour denoted the type of digit task: a black frame indicated sorting, while a red frame signalled a passive condition in which participants ignored the digits and pressed a button when the ‘fading square’ appeared. After a delay, participants recalled the initial image by selecting the memorised image. (B) The memory task stimulus set included two categories—tools and animals—each with 3 similarly-looking exemplars to increase task difficulty.

### Dual task

As displayed in Figure 1, panel A, on each trial, participants were asked to perform a memory task and a digit task (as an interrupter). A memory task trial started with a fixation cross (0.7° ×0.7°) displayed for 0.5s, followed by the memory target to be recalled after a delay period: an image (7° × 7°) of a morphed object (tool or animal), presented for 2 seconds.

After the memory target, a fixation cross (0.7° ×0.7°) was displayed on the screen for 0.75, 1, or 1.25 seconds. Then four squares (3°× 3° each, 1.5° width apart from each other) were presented centrally, each filled with one of the following colours, in this order, from left to right: blue, yellow, green, and red. The order of the colours was always the same and corresponded to the colours of the response button box. Within each square, a black digit was placed against a white background. The squares were presented for 1.5 seconds and were followed by a gray “fading” square (3°× 3°) gradually changing colour from gray to white to denote the time passing, displayed up to 1 second. Participants were instructed to mentally arrange the digits in ascending order and respond with the second digit in the ordered sequence when the “fading” square appeared.

To respond to the digit task, participants used a button box with four coloured buttons (blue, yellow, green, and red), which matched the colours of squares displayed on the screen with digits. Participants were required to press the button that matched the colour of the square associated with the digit they needed to respond to (the second item in a sorted list).

The digit task included three variations based on the level of task difficulty. In the easy digit task, the digits were randomly selected from between 1 and 4. In the hard digit task, the digits were selected from between 1 to 9. In this scenario, arranging a sequence of digits where each digit differs by more than one from the next would be more challenging within the restricted timeframe (hard digit task), compared to organising a sequence of digits where each one differs by exactly one from the next (easy digit task). The passive digit task served as a control and required only pressing the red button when the fading square appeared; participants were asked not to order the digits. In the passive task, the digits were specifically sampled the same way as for the hard digit task, so participants would not reorder the digits unintentionally. The type of the digit task was cued by the frame (thickness: 0.25°) around the squares with digits; the black frame informed about the active digit task – both hard and easy, and the red frame appeared for the passive digit task. Then, followed by another fixation cross displayed on the screen (0.7° ×0.7°) for 0.75, 1, or 1.25 seconds, participants were prompted to recall the image they had memorised. To test memory recall accuracy for the memorandum, four images were presented at the memory probe stage: one target image, one that was a different version of the same image, and two versions of a visually similar paired image (see Fig 1B). In a 4 AFC paradigm, participants selected the memorised image (7° ×7°) using the same button box, with each button mapped to the location of an image. After selecting an image, a 0.3° thick black frame appeared around it for 50 ms, followed by a blank screen for an inter-trial interval (jittered duration from 2 to 10 seconds).

Each participant underwent 4-5 training runs (9 trials each, lasting ∼2.5 minutes) to familiarise themselves with button assignments for memory recall and digit task responses, and a further 9 practice trials in the scanner before data collection. The number of digit task types was balanced in each training run.

The main experiment consisted of 8 runs, each run had 24 trials and lasted ∼6.5 minutes. The number of each type of digit task was balanced in each run (8 trials per type). The category of the memory target was also balanced in each run (4 tools and 4 animals were randomly chosen for each digit task type). The main experiment resulted in 192 trials in total, with 64 trials per digit task type.

### MR Data Acquisition

Participants were scanned on a 3T MRI scanner (Prisma Siemens) with a 32-channel head coil. During the scanning session, functional MRI scans were acquired with the following sequence parameters: 48 transversal slices collected in interleaved order with repetition time (TR) =1825 ms, echo time (TE) = 30 ms, multiband acceleration factor =2, voxel size = 2.5 × 2.5× 2.5 mm3, field-of-view = 192 × 192 mm, matrix size = 76 × 76, slice thickness = 2.5 mm, 0 mm slice gap, flip angle = 67°, covering the entire cerebrum except cerebellum. For each participant we also acquired a T1-weighted MR image (MP-RAGE; 176 sagittal slices, TR = 2300 ms, TE = 2.29 ms, voxel size = 1 × 1 × 1 mm3, field-of-view = 240 × 240 mm, matrix size = 240 × 240, slice thickness = 0.9 mm, 0 mm slice gap, TI = 900 ms, flip angle = 8°).

### Preprocessing

MRI data were pre-processed using SPM12 (Wellcome Department of Imaging Neuroscience, www.fil.ion.ucl.ac.uk/spm) in MATLAB 2021a (MathWorks, Inc., 2021). Functional MRI data were converted from DICOM to NIfTI format, spatially realigned to the mean functional scan and slice timing corrected. EPIs from the main experiment were smoothed (4 mm FWHM Gaussian kernel) to improve the signal-to-noise ratio, as applied in previous similar work (Jackson and Woolgar, 2018; Jackson et al., 2017). Structural scans were co-registered to the mean EPI and spatially normalised, using the segmentation and normalisation routines of SPM12, to derive the (inverse) normalisation parameters for individual ROI definition.

### ROI definition

Similar to previous studies investigating relevant over irrelevant representations in the MD network (Jackson and Woolgar, 2018; Jackson et al., 2017), we used the parcellated map of 13 frontal and parietal MD ROIs provided by Fedorenko et al., (2013; available online at imaging.mrc-cbu.cam.ac.uk/imaging/MDsystem). This definition of the MD network closely aligns with the previous definition by Duncan & Owen (2000), derived from meta-analytic data, and used in previous studies applying MVPA to MD regions (Jackson et al., 2017; Jackson & Woolgar, 2018; Jackson et al., 2021).

The MD ROIs included the left and right anterior inferior frontal sulcus (aIFS; centre of mass (COM) MNI coordinates = ±35 47 19, volume = 5.0 cm³), left and right posterior inferior frontal sulcus (pIFS; COM ±40 32 27, 5.7 cm³), left and right premotor cortex (PM; COM ±28 −2 56, 9.0 cm³), left and right inferior frontal junction (IFJ; COM ±44 4 32, 10.1 cm³), left and right anterior insula/frontal operculum (AIFO; COM ±34 19 2, 7.9 cm³), left and right intraparietal sulcus (IPS; COM ±29 −56 46, 34.0 cm³), and bilateral presupplementary motor area/anterior cingulate cortex (pre-SMA/ACC; COM 0 15 46, 18.6 cm³). We defined the early visual cortex (BA17: COM −1 −79 6, 31 cm3) from the Brodmann template provided with MRICroN (Rorden, 2007). All templates were provided in the MNI space and were then transformed to the native space by applying the inverse of the normalisation parameters for each participant using SPM12’s ‘segment’ and ‘normalise’ routine (Wellcome Department of Imaging Neuroscience, www.fil.ion.ucl.ac.uk/spm).

### General Linear Model (GLM)

We applied the General Linear Model (GLM) in SPM12 to estimate beta values for subsequent MVPA. For each run, we estimated separately the responses to each category (tool, animal) and digit task condition – easy, hard and passive. Two additional regressors representing “baseline” activity (tool, animal) during the inter-trial interval were also added to the model. This produced beta values for each run corresponding to 8 conditions. Trials were modelled as events as the delay period (i.e. starting at memory target offset until memory recall onset) and convolved with the canonical hemodynamic response of SPM12. In addition, motion parameters, and grand means of each run were also modelled as separate nuisance regressors resulting in fifteen regressors in total.

### Multivariate Pattern Analysis

To examine the fate of memory representations during the interrupter (digit) task, we used multivariate pattern analysis (MVPA) to decode the category of the maintained object (tool vs animal). Analyses were implemented in Python using scikit-learn (Pedregosa et al., 2011) and nilearn (Abraham et al., 2014).

For each participant, ROI, and interrupter condition (passive, easy, hard), delay-period activity patterns (GLM beta estimates, one per run × category) were extracted from each ROI mask using nilearn. A linear support vector machine (LinearSVC, C = 1) was trained to classify object category, with features standardised (z-scored) within the training set. We analysed the 13 multiple-demand (MD) regions (Fedorenko et al., 2013) and early visual cortex (V1) as an a priori sensory control.

Classification accuracy was estimated using leave-one-run-out cross-validation: on each of the eight folds, the classifier was trained on beta patterns from seven runs and tested on the held-out run, and accuracies were averaged across folds.

All group-level statistics were performed in Python (SciPy; Virtanen et al., 2020). To test whether category information was present, decoding accuracies were compared against chance (0.5) using one-tailed one-sample t-tests, for each ROI and condition. To assess the effect of the interrupter, we computed two planned contrasts per ROI — interrupter engagement ([easy + hard]/2 − passive) and interrupter demand (hard − easy) — each tested with a two-tailed paired t-test. For both the vs-chance and contrast analyses, p-values were corrected for multiple comparisons across the 13 MD ROIs using the Benjamini–Hochberg false-discovery-rate procedure (FDR); V1 was reported uncorrected as an a priori control. Effect sizes are reported as Cohen’s dz for within-subject comparisons.

### Cross-validated Mahalanobis (crossnobis) distance

As a complementary, noise-normalised measure of category information, we computed the cross-validated Mahalanobis (crossnobis) distance (Nili et al., 2014; Walther et al., 2016) between the tool and animal patterns, separately for each ROI and interrupter condition. Unlike decoding accuracy, which is bounded and conflates the magnitude of the pattern difference with the noise level, crossnobis provides a continuous, cross-validated estimate of representational distance that is unbiased under the null (expected value zero, and able to take negative values when no true difference is present).

Crossnobis distance was computed with custom scripts implementing the cross-validated Mahalanobis distance as described by Nili et al. (2014) and Walther et al. (2016), using Ledoit-Wolf shrinkage (Ledoit & Wolf, 2004) for noise covariance estimation. For each ROI, condition, and participant, distances were estimated using leave-one-run-out cross-validation. On each fold, the category difference vector (mean tool pattern − mean animal pattern) was computed separately in the training runs and in the held-out run, and the crossnobis distance was obtained as their inner product under a noise precision matrix: d = Δ_train ᵀ Σ⁻¹ Δ_test, averaged across the eight folds. Because the two difference vectors derive from independent partitions of the data, their noise terms are uncorrelated and cancel in expectation, yielding an unbiased distance estimate.

The noise precision matrix Σ⁻¹ was estimated from the residuals of the activity patterns around each condition × category cell mean (removing condition-level differences from the noise estimate), using Ledoit-Wolf shrinkage (Ledoit & Wolf, 2004), and was estimated from the training runs only on each fold and pooled across conditions, so that all conditions were expressed in common noise units. A single, consistent sign convention (tool − animal) was applied across folds and conditions.

Group-level statistics mirrored the decoding analysis. Category information was tested per ROI and condition against zero (one-tailed one-sample t-test). Interrupter effects were tested as two planned paired contrasts — engagement ([easy + hard]/2 − passive) and demand (hard − easy), two-tailed — corrected across the 13 MD ROIs using the Benjamini-Hochberg FDR procedure, with V1 reported uncorrected as an a priori control. As a validation of the estimator, we confirmed that a substantial proportion of distances were negative (as expected for an unbiased measure under the null).

### Behavioural Data Analysis

Behavioural responses were aggregated using a custom-written MATLAB script in Matlab 2021a (MathWorks, Inc., 2021) and analysed using Python Programming Language (v3.14, Van Rossum & Drake, 2009).

Although we ensured that each participant could perform the digit task with at least 60% accuracy prior to scanning, we also evaluated their performance during the experiment in the scanner. To ensure that participants attended to the digit task during memory maintenance, we calculated accuracy in each digit task condition and compared it against chance level (25% for active easy and hard; 50% for passive since the task presented a binary choice to press or not).

Three interrupter conditions were designed to create a demand gradient, with the passive condition least demanding, the easy condition more demanding, and the hard condition most demanding. To confirm that this manipulation was effective, we analysed digit-task performance (accuracy and reaction time), expecting poorer digit-task performance with increasing demand. We separately assessed the effect of interrupter demand on memory-task performance — recall accuracy and reaction time — and, to account for potential speed–accuracy trade-offs, additionally computed inverse efficiency scores (IES; Townsend & Ashby, 1983), defined as mean correct-trial reaction time divided by the proportion of correct responses for each participant and condition. IES summarises response cost across accuracy and speed, with higher values indicating poorer overall performance.

All responses were aggregated across runs within each participant, and analysed using repeated measures ANOVA with digit task condition as the main factor. Only correct responses entered response time analyses. Correct-trial reaction times below 0.2 s were treated as anticipatory and excluded from reaction-time analyses: 108 trials, 3.1% in memory task, and 220 trials, 6.3% in digit task. Post-hoc comparisons using Holm-Bonferroni correction were used to test the planned pairwise differences between conditions.

Two participants performed at chance levels across all digit-task conditions, indicating non-engagement, and were excluded only from digit-task analyses, resulting in twenty datasets. All twenty-two subjects’ datasets entered the memory task analysis.

### Bayesian analyses

To quantify evidence for the null hypothesis — of no difference between conditions (behavioural comparisons) and of no effect of the interrupter (neural contrasts) — we complemented all frequentist tests with Bayesian paired-samples t-tests. Bayes factors were computed using a Jeffreys–Zellner–Siow prior with the default scale factor r = 0.707 (Rouder et al., 2009), as implemented in Pingouin (Vallat, 2018). We report BF₀₁ (evidence for the null relative to the alternative) for comparisons expected to show no effect, and BF₁₀ (evidence for the presence of an effect) for tests of category information against the null (crossnobis vs. 0; decoding vs. chance). Bayes factors were interpreted using conventional benchmarks (anecdotal, 1–3; moderate, 3–10; strong, >10; Lee & Wagenmakers, 2013). For comparisons that violated normality (Shapiro–Wilk p < .05), Bayes factors were derived from the corresponding t-statistic and are therefore approximate.

### Network-level category information analyses

Because per-ROI tests may miss weak effects distributed across the network, we additionally tested both measures at the level of the MD network as a whole. For each participant and condition, decoding accuracies were averaged across the 13 MD ROIs. Crossnobis distances, which differ in scale between ROIs (e.g., distances in IPS were an order of magnitude larger than in frontal ROIs), were first divided by each ROI’s standard deviation across participants and conditions, so that each ROI contributed equally while condition differences within an ROI were preserved, and then averaged across ROIs.

Network means were tested against the null (0.5 for decoding; 0 for crossnobis) in each condition with two-tailed one-sample t-tests, and the engagement and demand contrasts were tested with two-tailed paired t-tests, complemented by JZS Bayes factors, Wilcoxon signed-rank tests and sign-flip permutation tests (100,000 permutations). Robustness was assessed by recomputing the network contrasts after leaving out each ROI in turn and, for crossnobis, by repeating the analysis on raw and voxel-normalised distances (distance divided by the number of voxels) and after excluding bilateral IPS. Results were confirmed with linear mixed-effects models fitted to all ROIs (statsmodels; Seabold & Perktold, 2010), with fixed effects of the two contrasts and of ROI, and by-participant random intercepts and random slopes for both contrasts; a likelihood-ratio test of the contrast × ROI interaction assessed whether interrupter effects differed between ROIs. Finally, to assess agreement between the two measures, we correlated participants’ network-level contrasts for decoding and crossnobis.

## Results

### Performance on the digit task reflected interrupter demand manipulation

First, the difficulty manipulation of the interrupting digit task was confirmed by reaction times, which showed a strong graded effect of demand (F(2, 38) = 28.89, p < .001, partial η² = .60). Responses were slowest in the hard condition (M = 0.50 s, SD = 0.11 s), intermediate in the easy condition (M = 0.42 s, SD = 0.09 s), and fastest in the passive condition (M = 0.38 s, SD = 0.11 s). The hard condition was significantly slower than both easy (t(19) = 8.12, p < .001, d = 1.82, BF₁₀ > 1000) and passive (t(19) = 5.77, p < .001, d = 1.29, BF₁₀ > 1000), and easy was slower than passive (t(19) = 2.40, p = .027, d = 0.54, BF₁₀ = 2.28; all Holm-corrected).

Accuracy provided converging evidence. Although the omnibus effect of condition was not significant (F(2, 38) = 1.60, p = .215), the critical contrast between the hard and easy conditions was reliable: accuracy was lower in the hard condition (M = 87.5%, SD = 11.7%) than in the easy condition (M = 93.6%, SD = 6.8%; t(19) = 3.60, p = .006, Holm-corrected; d = 0.81, BF₁₀ = 20.92). Passive accuracy (M = 90.2%, SD = 16.5%) is reported for completeness but was not treated as a graded-difficulty baseline, as it reflects a qualitatively different, single-response task. Together, the reaction-time and accuracy results confirm that the difficulty manipulation was successful.

### Effect of Interrupter and its Demand on Memory Recall

Memory recall accuracy reflected the cost of interruption (F(2, 42) = 4.16, p = .023, partial η² = .17; Figure 2A). Relative to the passive condition (M = 85.2%, SD = 9.0%), accuracy was reduced when a hard interrupter was performed during the delay (M = 81.1%, SD = 10.9%; t(21) = 2.80, p = .033, Holm-corrected; dz = 0.60, BF₁₀ = 4.65). A comparable numerical reduction in the easy condition (M = 80.8%, SD = 10.3%; t(21) = 2.27, raw p = .034, Holm p = .068, BF₁₀ = 1.81) was inconclusive after correction. Critically, accuracy did not differ between the easy and hard conditions, with moderate evidence for the null (Wilcoxon signed-rank, p = .52; BF₀₁ = 4.43).

**Figure 2.**
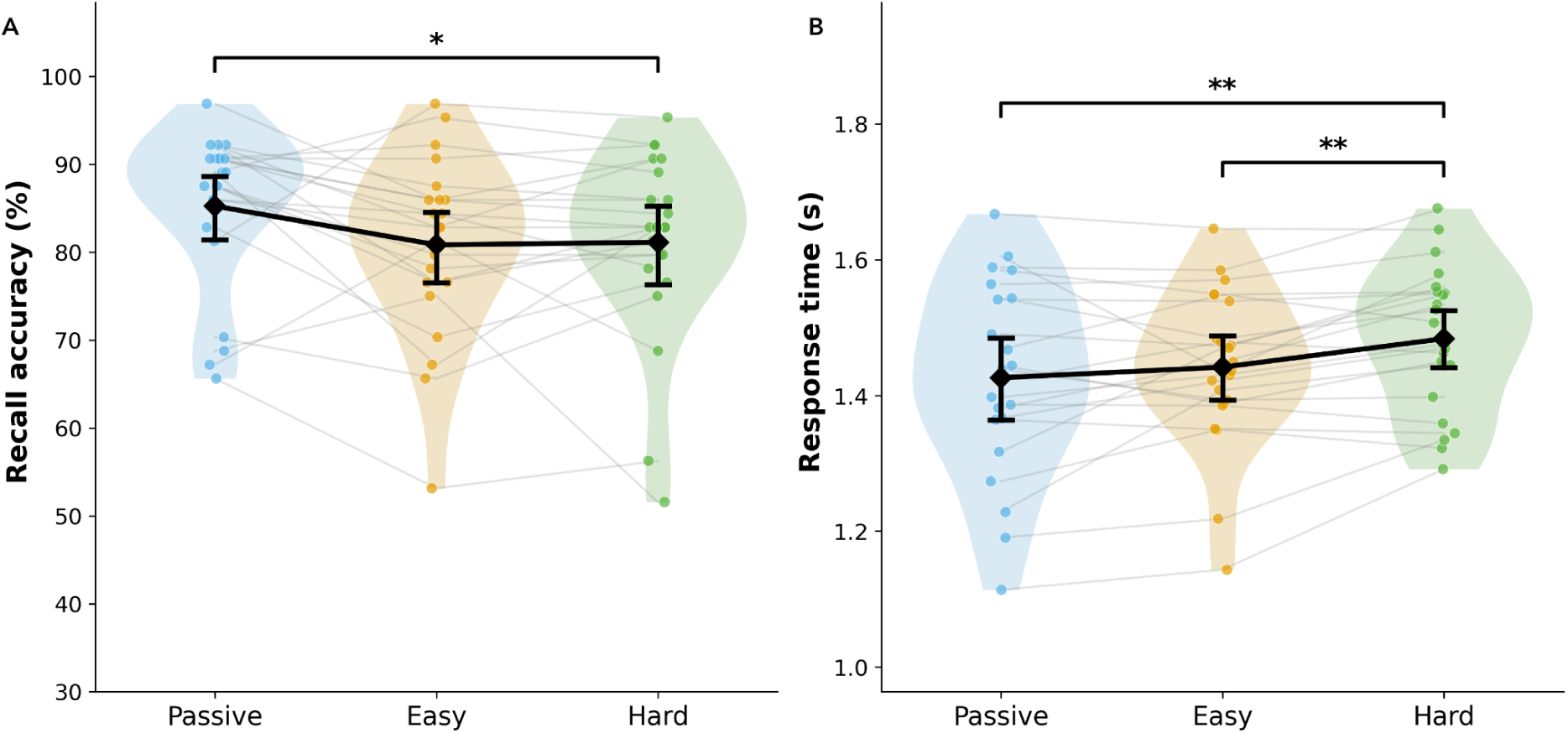
Effects of digit task difficulty on WM recall accuracy and response times. Memory recall performance across interrupter conditions (N = 22). (A) Recall accuracy was reduced by the engagement with the hard interrupter in comparison to the passive (p = .033, Holm-corrected), with no difference between easy and hard. (B) Response times were slower in the hard condition than in both the easy and passive conditions (p < .05, Holm-corrected). Diamonds show group means; error bars show 95% confidence intervals; gray lines show individual participants. *p < .05, **p ≤ .01.

Response times showed a complementary, graded pattern (F(2, 42) = 7.39, p = .002, partial η² = .26; Figure 2B). Recall was slower in the hard condition (M = 1.48 s, SD = 0.11 s) than in both the passive (M = 1.43 s, SD = 0.14 s; t(21) = 3.16, p = .010, Holm-corrected; dz = 0.67, BF₁₀ = 9.37) and easy (M = 1.44 s, SD = 0.12 s; t(21) = 3.60, p = .005, dz = 0.77, BF₁₀ = 22.81) conditions. The easy and passive conditions did not differ (t(21) = 0.98, p = .34; BF₀₁ = 2.93).

To capture response cost across both accuracy and speed, we examined inverse efficiency scores (IES). IES differed across conditions (F(2, 42) = 5.41, p = .008, partial η² = .20). Specifically, IES was higher (poorer performance) in the hard than the passive condition (M = 1.87, SD = 0.37 vs. M = 1.70, SD = 0.33; t(21) = 3.29, p = .010, Holm-corrected; dz = 0.70, BF₁₀ = 12.20), whereas the easy (M = 1.82, SD = 0.32) and hard conditions did not differ (Wilcoxon signed-rank, p = .41; BF₀₁ = 2.88), nor did easy and passive after correction (t(21) = 2.22, raw p = .037, Holm p = .073, BF₁₀ = 1.72).

### Decoding of memory content across the multiple-demand network and early visual cortex

We first asked whether the maintained category (tool vs. animal) could be decoded from delay-period activity (Figure 3, left). Decoding was reliably above chance in visual cortex in every condition (V1: accuracy 0.58–0.60, dz = 0.94–1.35, all p < .001; all BF₁₀ > 100). Within the MD network, category information was carried most consistently by bilateral IPS: IPS-R decoded above chance in all three conditions (dz = 0.59–0.76, p ≤ .039, BF₁₀ = 4.3–21.9) and IPS-L in the passive and hard conditions (dz = 0.72–0.74, p ≤ .019, BF₁₀ = 14.2–17.2), with a comparable trend in the easy condition (dz = 0.48, p = .056, BF₁₀ = 1.8). Beyond IPS, above-chance decoding was sparse and condition-specific: no other region was significant in more than one condition (passive: IFJ-R, pIFS-R, aIFS-R; easy: IFJ-L, AIFO-L; hard: pIFS-L). In total, 5, 3, and 3 of 13 MD regions decoded above chance in the passive, easy, and hard conditions, respectively; five core regions (ACC, bilateral PM, left aIFS, right AIFO) decoded above chance in no condition. Mean decoding across the MD network (0.53) was lower than in the visual cortex (0.59).

**Figure 3.**
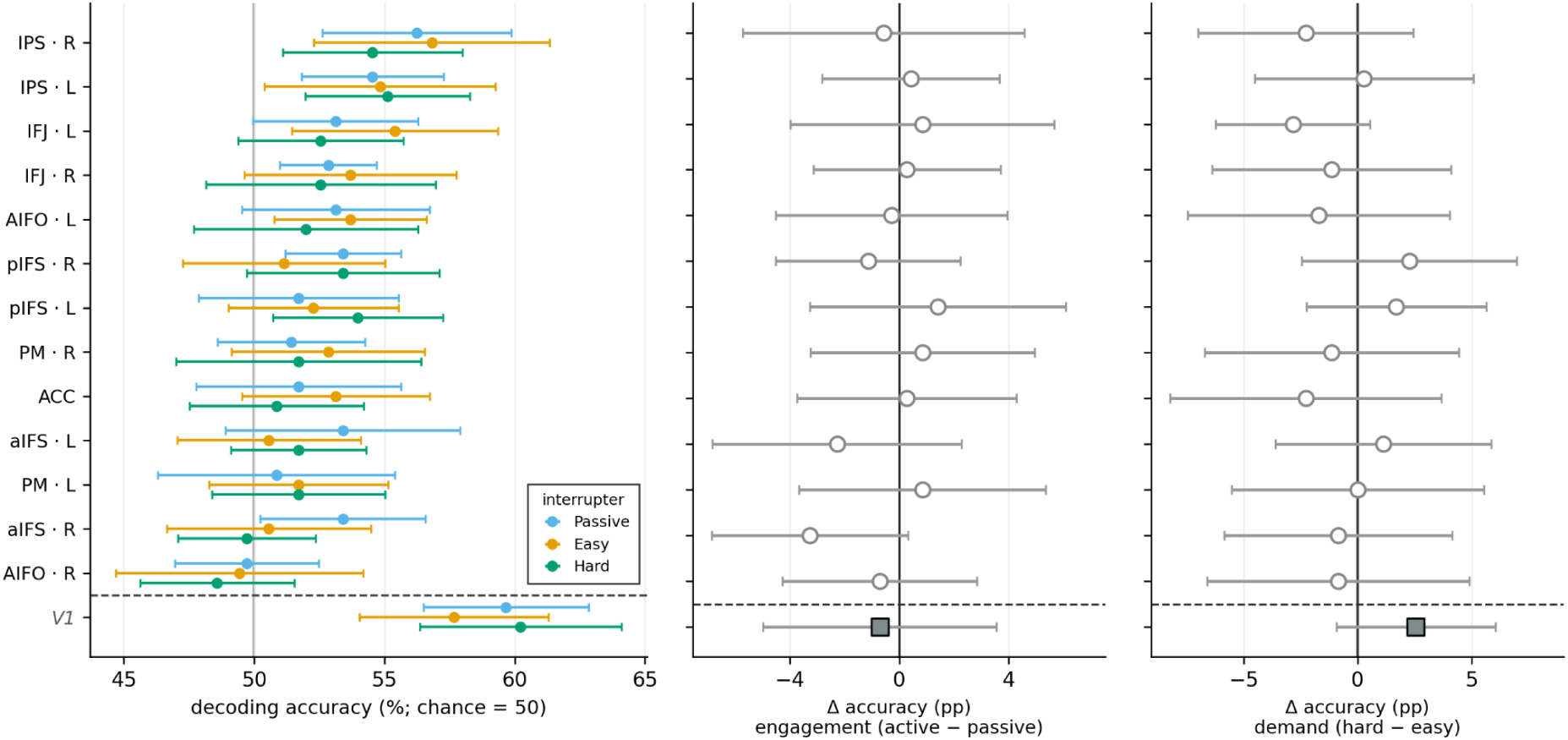
Category information is present in the multiple-demand network but not modulated by interrupter engagement or demand in individual regions. Decoding of the maintained category (tool vs. animal) from delay-period activity across 13 multiple-demand (MD) regions in N = 22 participants; bilateral early visual cortex (V1) is included as a sensory control below the dashed line. MD regions are ordered by mean decoding accuracy. (Left) Decoding accuracy for each interrupter condition, expressed as distance from chance (0.5, vertical line); points denote group means ± 95% CI. Within the MD network, category information was carried most consistently by bilateral IPS. (Middle) No interrupter task effect (active − passive) and (Right) interrupter demand effect (hard − easy) were found in any of the ROIs; points show mean paired differences ± 95% CI, with the vertical line at zero. Neither contrast was significant in any MD region after FDR correction; V1 is reported uncorrected (gray square).

### Category Decoding is not modulated by interrupter engagement or demand

As illustrated in Figure 3, neither interrupter engagement (Middle), nor demand (Right) significantly altered decoding accuracy in any MD region after FDR correction (both contrasts: 0/13 regions). Bayesian paired t-tests provided moderate evidence for the null in most regions (engagement: 11 of 13; demand: 10 of 13). For the task-engagement contrast, evidence favoured no effect throughout, including in bilateral IPS (IPS-L: Δ = 0.004, 95% CI [−0.028, 0.037], t(21) = 0.27, p = .787, BF₀₁ = 4.33; IPS-R: Δ = −0.006, [−0.057, 0.046], t(21) = −0.23, p = .821, BF₀₁ = 4.38) and V1 (BF₀₁ = 4.25). For the demand contrast, evidence again favoured the null in IPS-L (Δ = 0.003, t(21) = 0.12, p = .903, BF₀₁ = 4.45) and, more weakly, IPS-R (Δ = −0.023, p = .329, BF₀₁ = 2.87) and V1 (BF₀₁ = 1.64). Thus, category information was decodable during the delay, and no individual region showed a reliable effect of either the engagement or the difficulty of the interrupter (for network-level tests, see Network-level category information analyses).

### Category representations are maintained across the multiple-demand network and early visual cortex

Category information (crossnobis distance vs. 0) was reliably represented in the visual cortex and across much of the multiple-demand (MD) network as shown in Figure 4 (Left). Distances were largest and most robust in V1, in every condition (dz = 1.60–1.70, all p < .001; all BF₁₀ > 73,000). Within the MD network, category information was strongest in bilateral intraparietal sulcus (IPS-L: dz = 0.94–1.49; IPS-R: dz = 0.89–1.31; all p ≤ .002; all BF₁₀ > 72). Six regions carried reliable information under all three interrupter levels — bilateral IPS, inferior frontal junction (IFJ-L, IFJ-R), and posterior inferior frontal sulcus (pIFS-L, pIFS-R) — while others reached significance under some conditions only. Overall, 10, 9, and 6 of 13 MD regions were significant in the passive, easy, and hard conditions, respectively; only right AIFO failed to reach significance throughout.

**Figure 4.**
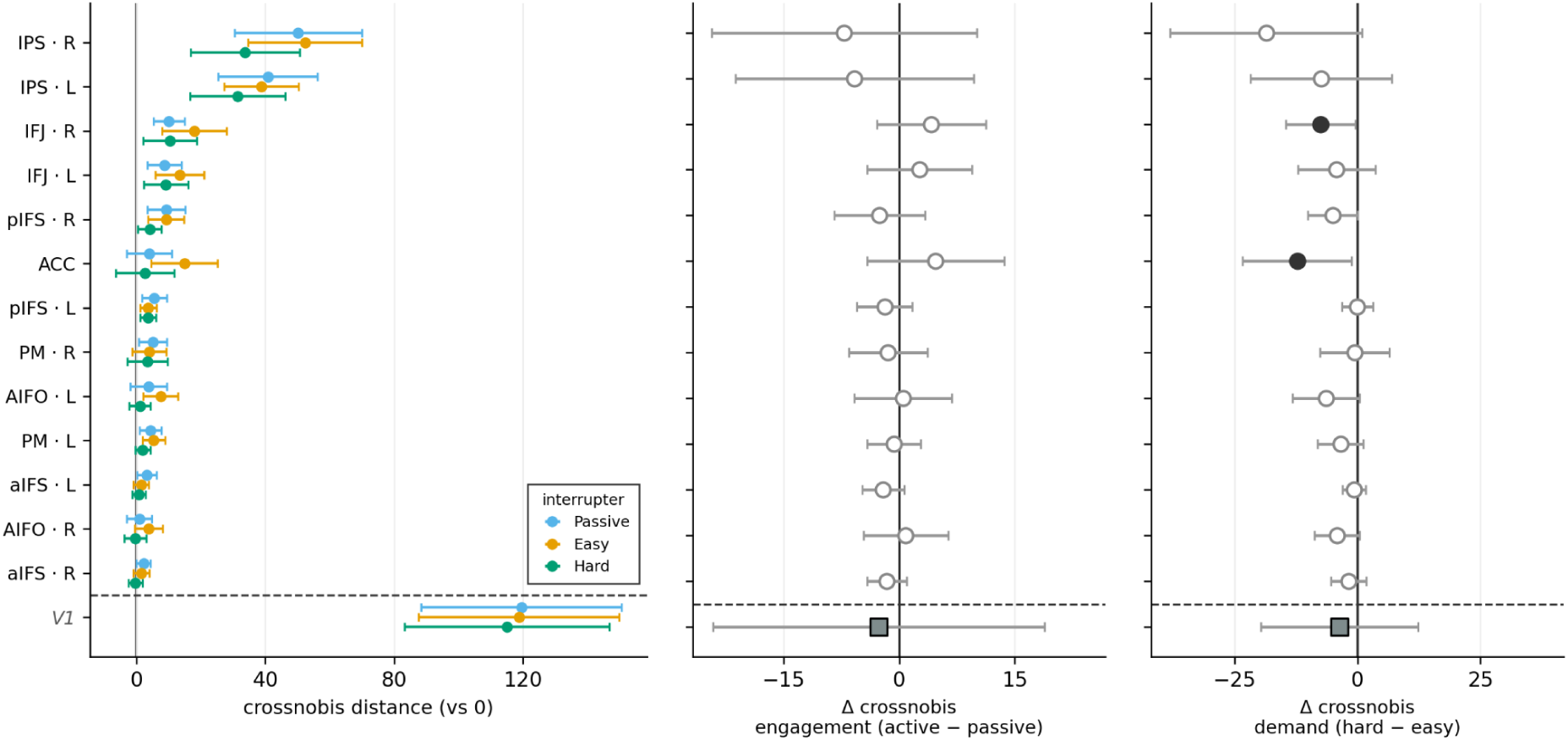
Category representations in the multiple-demand network are present but not reliably modulated by interrupter engagement or demand in individual regions. Cross-validated Mahalanobis (crossnobis) distance between the tool and animal patterns during the delay period, across 13 multiple-demand (MD) regions, in N = 22 participants; early visual cortex (V1) is included as a sensory control (shown below the dashed line). MD regions are ordered by mean distance. (Left) Crossnobis distance in each interrupter condition (passive, easy, hard; colours), relative to zero (the null for this cross-validated measure; vertical line). Points show the group means ± 95% CI. Within the MD network, category information was strongest and most consistent in bilateral IPS, and was present across much of the network. (Middle) Engagement effect of the interrupter task ([easy + hard]/2 − passive) and (Right) its demand (hard − easy); points show the mean paired difference ± 95% CI, with the vertical line at zero. Neither contrast was significant in any MD region after FDR correction; filled circles indicate a 95% CI excluding zero (uncorrected). Distances were tested against zero (one-tailed one-sample t-tests) and contrasts with two-tailed paired t-tests, FDR-corrected across the 13 MD ROIs; V1 is reported uncorrected.

### Interruption engagement and demand do not reliably modulate representational distance in individual regions

As illustrated on Figure 4, neither the engagement (Middle) with the interrupter nor its demand (Right) modulated crossnobis distance in any MD region after FDR correction (both contrasts: 0/13 regions). For the task-engagement contrast, Bayesian paired t-tests favoured the null in all regions (moderately in eight and anecdotally in five), including bilateral IPS (IPS-L: Δ = −5.8, 95% CI [−21.3, 9.7], t(21) = −0.78, p = .445, BF₀₁ = 3.41; IPS-R: Δ = −7.1, [−24.3, 10.1], t(21) = −0.86, p = .398, BF₀₁ = 3.21) and V1 (Δ = −2.6, p = .802, BF₀₁ = 4.36). For the demand contrast, all regional effects were negative in direction — a tendency toward lower discriminability under the hardest interrupter — but none survived correction. Bayesian evidence was mixed: it favoured the null moderately in three regions (PM-R, aIFS-L, pIFS-L) and anecdotally in four (e.g., IPS-L: Δ = −7.4, [−21.8, 7.0], t(21) = −1.07, p = .298, BF₀₁ = 2.71), and was inconclusive in the remaining six (BF₀₁ = 0.52–1.00; e.g., IPS-R: Δ = −18.6, [−38.2, 0.9], t(21) = −1.98, p = .060, BF₀₁ = 0.86); at uncorrected thresholds, IFJ-R and ACC showed reductions. In V1, evidence favoured the null (BF₀₁ = 4.04). Because the demand effects were uniformly negative but individually weak, we additionally tested them at the network level (see Network-level category information analyses).

## Network-level category information results

Per-ROI tests may miss weak effects distributed across the network: for crossnobis, all 13 regional demand effects were negative in direction, yet none survived correction. We therefore tested both measures at the level of the MD network as a whole (Figure 5).

**Figure 5.**
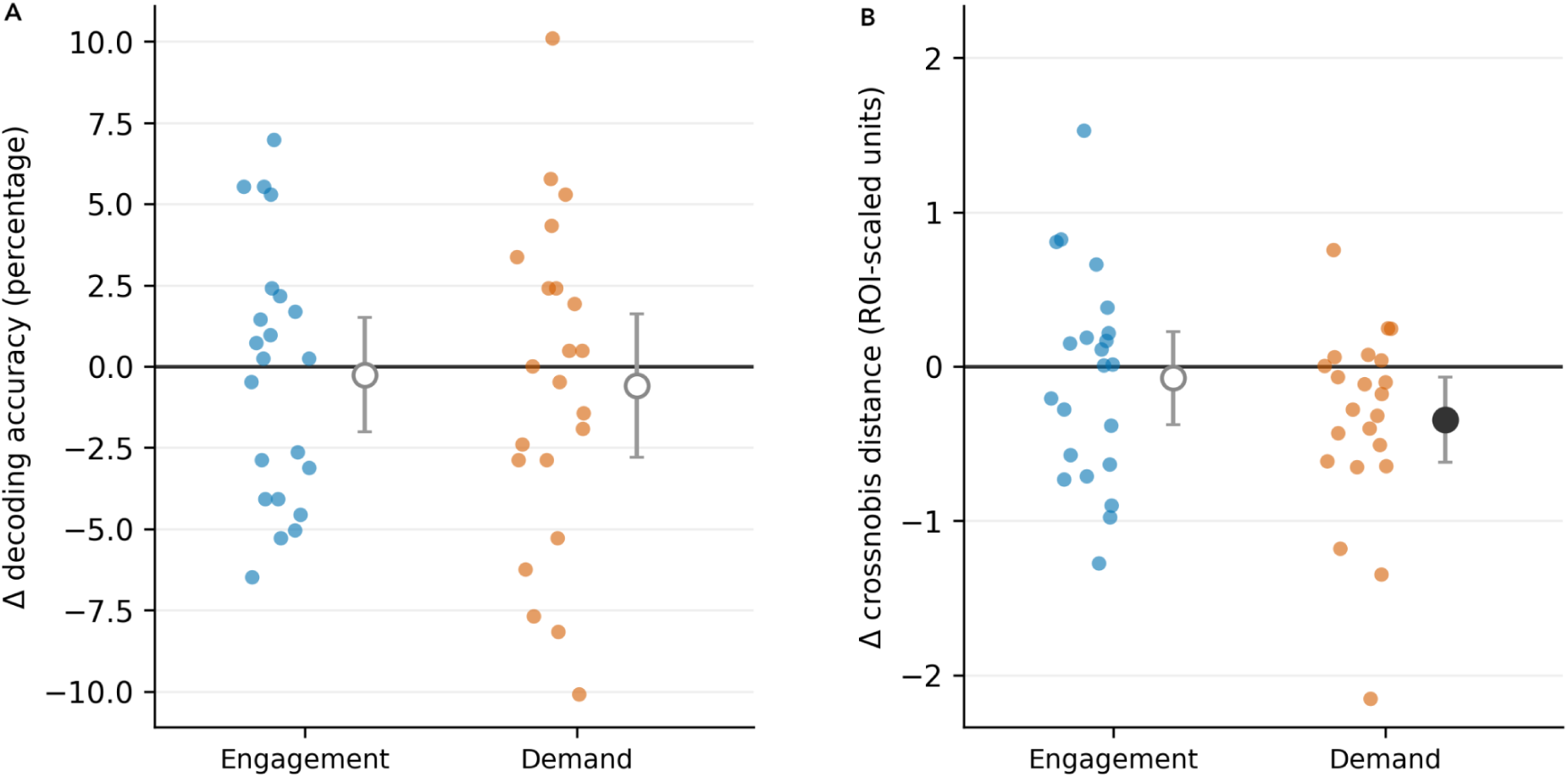
Network-level category information. Each participant’s MD-network contrasts for interrupter engagement ( [easy + hard]/2 − passive) and interrupter demand (hard − easy), for (A) decoding accuracy (percentage) and (B) crossnobis distance (ROI-scaled units; each ROI divided by its standard deviation before averaging across the 13 MD ROIs), in N = 22 participants. Coloured points show individual participants; circles and error bars show group means ± 95% CI, with the horizontal line at zero. Filled circles indicate a 95% CI excluding zero.

Category information was present in the MD network in every condition, for both decoding accuracy (passive: t(21) = 3.99, p < .001; easy: t(21) = 3.10, p = .005; hard: t(21) = 2.98, p = .007) and crossnobis distance (passive: t(21) = 5.12, p < .001; easy: t(21) = 5.07, p < .001; hard: t(21) = 2.80, p = .011).

The two measures converged for interrupter engagement: neither decoding accuracy (t(21) = −0.30, p = .77, dz = −0.06, BF₀₁ = 4.31) nor crossnobis distance (t(21) = −0.51, p = .61, dz = −0.11, BF₀₁ = 3.98) was affected by the interrupter, with moderate Bayesian evidence for the null in both cases (Figure 5).

For interrupter demand, the measures diverged. Crossnobis distance was lower in the hard than in the easy condition across the network (t(21) = −2.62, p = .016, dz = −0.56, BF₁₀ = 3.36; Figure 5B). This effect was not carried by any single region: it remained significant when each ROI was left out in turn (p = .012–.023), when bilateral IPS was excluded (p = .019), and for raw and voxel-normalised distances (p = .011–.014), and it was confirmed by Wilcoxon signed-rank and sign-flip permutation tests (both p = .010). Decoding accuracy, in contrast, showed no effect of demand (t(21) = −0.55, p = .59, dz = −0.12, BF₀₁ = 3.90; leave-one-ROI-out: p = .45–.72; Figure 5A).

Mixed-effects models across all MD ROIs confirmed this pattern. For crossnobis, demand reduced category information (b = −0.34, 95% CI [−0.60, −0.09], p = .009), whereas engagement had no effect (b = −0.07, [−0.36, 0.21], p = .61). For decoding, neither contrast was significant (demand: b = −0.006, [−0.027, 0.015], p = .58; engagement: b = −0.003, [−0.019, 0.014], p = .77). For neither measure did interrupter effects differ between ROIs (crossnobis: χ²(24) = 19.02, p = .75; decoding: χ²(24) = 11.89, p = .98), indicating a distributed rather than region-specific pattern.

Despite this divergence at the group level, the two measures were related across participants: network-level contrasts correlated between decoding and crossnobis for both engagement (r = .77, p < .001) and demand (r = .49, p = .022). Thus, the demand effect appears to be present in both measures but is detectable only with crossnobis distance, a continuous measure that retains information about the magnitude of pattern differences, and not with classification accuracy, which is coarse (Walther et al., 2016).

## Discussion

The present study examined how an interrupting task, and its parametrically varied demand, affects the neural representation of maintained content in the multiple-demand (MD) network. Using two complementary multivariate measures—multivariate pattern classification and cross-validated representational distance (crossnobis)—we found that the category of the maintained object was reliably represented during the delay and remained largely unaffected by the interrupter: its engagement did not alter category information in any MD region or at the network level, according to either measure, and its increasing difficulty did not alter category information in any individual region, producing only a modest network-wide reduction in crossnobis distance that was not evident in decoding accuracy. Behaviour, by contrast, was disrupted. Recall accuracy was reduced by the engagement with an interrupter, irrespective of its difficulty, whereas reaction times were slowed only by the most demanding interrupter. Thus, interruption impaired behavioural performance while leaving the categorical representation of maintained content across the MD network and early visual cortex largely intact, suggesting that the behavioural consequences of interruption are not necessarily reflected in this aspect of the memory representation.

What happens to a maintained memory when another task (such as interrupter) competes for cognitive resources? We considered three accounts of how the MD network might respond to such interference, each predicting a distinct relationship between category information and interrupter demand. Our results are most consistent with demand-insensitive maintenance, qualified by a weak, distributed cost of demand. An active-protection account would predict that category information is preserved or enhanced as demand increases, consistent with the demand-dependent enhancement of task-relevant codes previously observed in this network (Woolgar et al., 2011; Jackson et al., 2021). We found no such enhancement in any region or with either measure. Conversely, competition for shared representational resources would predict a decline in category information with increasing demand. Here, the evidence depended on the scale and the measure: no individual region showed a reliable reduction, but crossnobis distance decreased modestly across the MD network under the hardest interrupter, an effect that was not detectable in decoding accuracy. Interrupter engagement, by contrast, left category information unchanged on both measures, with moderate Bayesian evidence for the null. Category information thus remained detectable in every condition, consistent with a representation that is maintained largely independently of the interrupting task, with at most a weak, distributed loss of discriminability when that task becomes most demanding — a pattern compatible with modest competition for shared resources that leaves the format of the code intact.

This finding fits the characterisation of the MD network as an adaptive coder that preferentially represents information relevant to the task at hand, rather than all information available (Duncan, 2001, 2010; Jackson & Woolgar, 2018). Two features of our data support this interpretation. First, category information was distributed unevenly across the brain: it was strongest in the visual cortex and, within the MD network, in the intraparietal sulcus (IPS), the MD region closest to the sensory hierarchy, and weaker across the remaining MD network — sparse when measured with decoding accuracy, but detectable in most regions with crossnobis distance. This gradient converges with recent findings from Shashidhara et al. (2024), who compared multivariate patterns evoked by faces and buildings across three executive tasks. Whereas task identity was strongly discriminated within core MD regions, stimulus category was weakly represented within those regions. Category information was instead strongest in the visual cortex and in parietal and frontal regions adjacent to the MD core, which may relay stimulus information into the network (Shashidhara et al., 2024). Second, the memorandum’s category was not modulated by interrupter engagement, and only weakly by its demand. Because category was relevant to the trial as a whole but incidental to the interrupting task performed during the delay, it was not differentially prioritised as the demands of the ongoing task increased. Instead, its representation remained largely stable. On this account, the stability we observed need not reflect a dedicated protective mechanism, but may instead arise from a general property of adaptive coding: information that is not currently being prioritised can remain represented without being dynamically altered as task demands change.

Critically, the stability of the category code dissociated from a reliable behavioural cost: the engagement with an interrupter reduced recall accuracy, while increasing interrupter demand slowed responses. The behavioural cost therefore is unlikely to be attributed to a loss of category-level information in the MD network. The modest network-wide reduction under the hardest interrupter coincided with slower recall, but its small size, and its absence in decoding accuracy, make it an unlikely sole source of the behavioural cost; in particular, the accuracy cost of interruption occurred without any measurable change in category information. Earlier work has shown that interruption can disrupt frontal–sensory connectivity, which is reinstated once the interrupter has passed (Clapp et al., 2010). Our results suggest that such disruption need not entail loss of the maintained representation itself. Instead, the cost may arise at a finer representational scale (Brady & Alvarez, 2011; Hallenbeck et al., 2021), for example as a loss of precision for information outside the focus of attention (LaRocque et al., 2015), or at later stages of retrieval and attentional selection, when the memory must be re-selected and prepared for use (Lewis-Peacock et al., 2012; Myers et al., 2017). Our memory probe required participants to identify the target among four perceptually similar images from the same category, such that accurate performance depended on item-specific information that category-level representations alone could not support.

This account makes a testable prediction: exemplar-level representational similarity analysis should reveal a reduction in item-specific information under interruption despite preserved category information, such that the broader categorical representation remains intact while finer-grained detail is degraded. Testing this dissociation would provide a direct test of whether interruption selectively affects mnemonic detail. A complementary causal approach would be to disrupt MD regions during the delay using TMS. Concurrent TMS–fMRI has shown that dorsolateral prefrontal cortex causally shapes the coding of task-relevant information (Jackson et al., 2021). Applied during interruption, this approach could establish whether MD activity is necessary for maintaining the memorandum, for performing the interrupting task, or for coordinating the two.

Finally, the network-level reduction in category information under demand was detected by crossnobis distance but not by decoding accuracy, and was modest in size. Although the two measures were correlated across participants, this effect should be treated as preliminary until replicated, ideally with designs that sample interrupter demand more finely and with more trials per condition.

In conclusion, we found that the category of a maintained object remained reliably represented across the MD network during interruption, and was at most modestly weakened as the interrupting task became more demanding, despite a concurrent decline in behavioural performance. Thus, the behavioural cost of interruption was not reflected in a loss of the categorical representation of maintained content, suggesting that its effects may emerge at a finer representational or control level. Future work should test whether item-specific information is more vulnerable to interruption and establish the causal contribution of MD regions to memory maintenance during interruption, potentially informing approaches to enhancing cognitive resilience in clinical populations.

